# Enhanced Place Specificity of the Parallel Auditory Brainstem Response: An Electrophysiological Study

**DOI:** 10.1101/2024.03.10.584313

**Authors:** Thomas J Stoll, Ross K Maddox

## Abstract

**Purpose:** This study investigates the effect of parallel stimulus presentation on the place specificity of the auditory brainstem response (ABR) in human listeners. Frequency-specific stimuli do not guarantee a response from the place on the cochlea corresponding only to that characteristic frequency – especially for brief and high-level stimuli. Adding masking noise yields responses that are more place specific, and a prior modeling study has suggested similar effects when multiple frequency-specific stimuli are presented in parallel. We tested this hypothesis experimentally here, comparing the place specificity of responses to serial and parallel stimuli at two stimulus frequencies and three stimulus rates.

**Methods:** Parallel ABR (pABR) stimuli were presented alongside high-pass filtered noise with a varied cutoff frequency. Serial presentation was also tested by isolating and presenting single-frequency stimulus trains from the pABR ensemble. Latencies of the ABRs were examined to assess place specificity of responses. Response bands were derived by subtracting responses from different high pass noise conditions. The response amplitude from each derived response band was then used to determine how much individual frequency regions of the auditory system were contributing to the overall response.

**Results:** We found that parallel presentation improves place specificity of ABRs for the lower stimulus frequency and at higher stimulus rates. At a higher stimulus frequency, serial and parallel presentation were equally place specific.

**Conclusion:** Parallel presentation can provide more place specific responses than serial for lower stimulus frequencies. The improvement increases with higher stimulus rates and is in addition to the pABR’s primary benefit of faster test times.

## Introduction

Although the cochlea is organized tonotopically, frequency-specific stimuli can elicit broad excitation patterns when presented at high levels [1–4]. ‘Place specificity’ is a way of describing this excitation pattern, where poor place specificity indicates a spread of excitation to characteristic frequencies (CF) beyond the frequency spectrum of the stimulus. Frequency-specific stimuli at low to moderate levels elicit place-specific responses, but place specificity worsens at the higher levels which may be needed to evoke a response from an ear with elevated audiometric thresholds. This decrease in place specificity results in responses from off-frequency regions, potentially producing inaccurate results or missed diagnoses [5, 6].

Place specificity can be improved by presenting masking noise alongside test stimuli [1, 7–10]. Masking noise is typically constructed such that the noise has power in the frequency regions where a response is not desired. The noise continuously and randomly excites the masked regions, and the adaptation and refractoriness of the excitable cells prevents these regions from producing responses which are time-locked to the stimulus [4]. Thus, the contributions of the masked regions to the average response are diminished. Despite the benefit of improved place specificity provided by the addition of masking noise, it is rarely used in practice because it reduces response size and modifies response morphology, therefore increasing test times (the limiting factor for ABR exams) and making interpretation difficult [11, 12].

Frequency-specific ABR exams are often used to estimate hearing thresholds across a range of frequencies for individuals who are not able to provide a behavioral response [1, 8]. This is accomplished by presenting short tonebursts of different frequencies to each ear while recording brain signals using electroencephalography (EEG) and averaging many (typically thousands) of trials, resulting in a stereotyped waveform consisting of several peaks. Standard ABR exams test one ear and one frequency at a time, and the stimulus intensity is varied until a threshold level is found. Clinically, ABR exams are most often used for early diagnosis of hearing loss in infants, and the test must be carried out while they are asleep to prevent motion artifact, which limits recording time. This may lead to incomplete results, forcing the clinician to infer thresholds from incomplete information, or ask patients to return for an additional visit [12].

Our lab developed the parallel ABR (pABR) paradigm to reduce test times [13]. This reduction is accomplished by utilizing randomized stimulus timing to test multiple frequencies in both ears simultaneously. Due to the random stimulus timing, responses to different frequencies are statistically independent and can be extracted from a single EEG recording. We further hypothesized the simultaneous presentation could improve place specificity of responses with no additional masking noise, as each stimulus may serve as both test stimulus and masker since the toneburst trains are uncorrelated (Fig. 1).

**Fig. 1.**
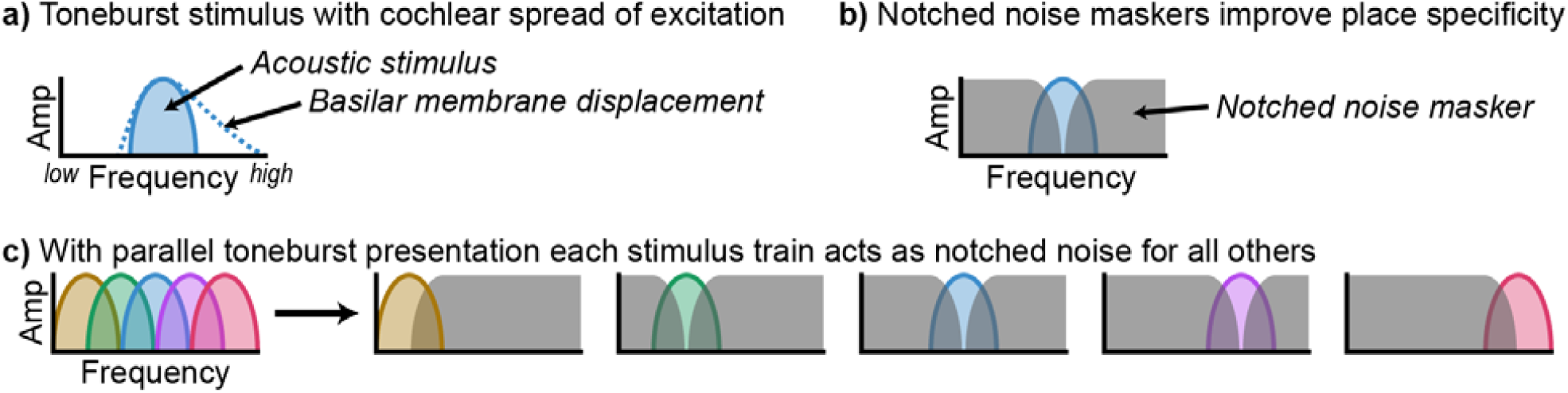
Cartoon representation of the hypothesis that parallel presentation improves place specificity. (a) A stimulus of one frequency is shown in the frequency domain. The solid line and filled area represent the frequency spectrum of the stimulus, while the dotted line represents the asymmetric spread of excitation on the cochlea. (b) Notched noise is added to the single frequency stimulus, preventing the spread of excitation to regions beyond the stimulus frequency. (c) Multiple frequency-specific stimuli are presented at the same time. When computing the response to one frequency, the stimuli of other frequencies act as masking noise. From Polonenko & Maddox [13], licensed CC BY-NC.

We previously used computational modeling to examine pABR response spread and found that parallel presentation produces responses which are more place specific than standard methods [14]. We used two models of the auditory periphery to simulate responses across a range of stimulus rates and levels for both parallel (all toneburst trains summed and presented together) and serial (a toneburst train of one frequency presented in isolation) stimuli. Both models suggest parallel presentation is as place specific as serial presentation at low levels and more place specific than serial presentation at high levels. This improvement was greater at higher stimulus rates and most pronounced for lower stimulus test frequencies. These models, however, generate auditory nerve responses, which correspond to wave I of the ABR, while the more robust and thus clinically relevant response component is wave V, which is generated by a mixture of midbrain sources including the lateral lemniscus and inferior colliculus [1].

In this study, we used a derived-band stimulus and analysis paradigm while presenting pABR stimuli either in parallel or serially to test the place specificity of responses in human subjects. The derived-band paradigm uses masking noise to isolate responses to specific frequency regions of the auditory system [15, 16]. High-passed noise (HPN) is used to prevent regions of the cochlea with a characteristic frequency higher than the HPN filter cutoff from contributing to the average responses. This improvement in cochlear place specificity should then be inherited by later stages in the auditory pathway. By varying this cutoff frequency, multiple responses with overlapping response bands can be collected. These responses are then subtracted to examine the responses generated by specific frequency regions (Fig. 2). Due to the reduced signal-to-noise ratio (SNR) of derived responses, we examined both un-subtracted and derived responses in this paper. Derived responses have been examined previously for serial ABRs [15, 16] and auditory steady-state responses [17] to show both are place specific. The methods used in this study, which are closely based on these previous studies, investigate the place specificity of the pABR and compare it with standard ABRs. We find that the pABR is place specific in general, with improved place specificity over serial ABRs at a low stimulus frequency, and no differences at a higher stimulus frequency.

**Fig. 2.**
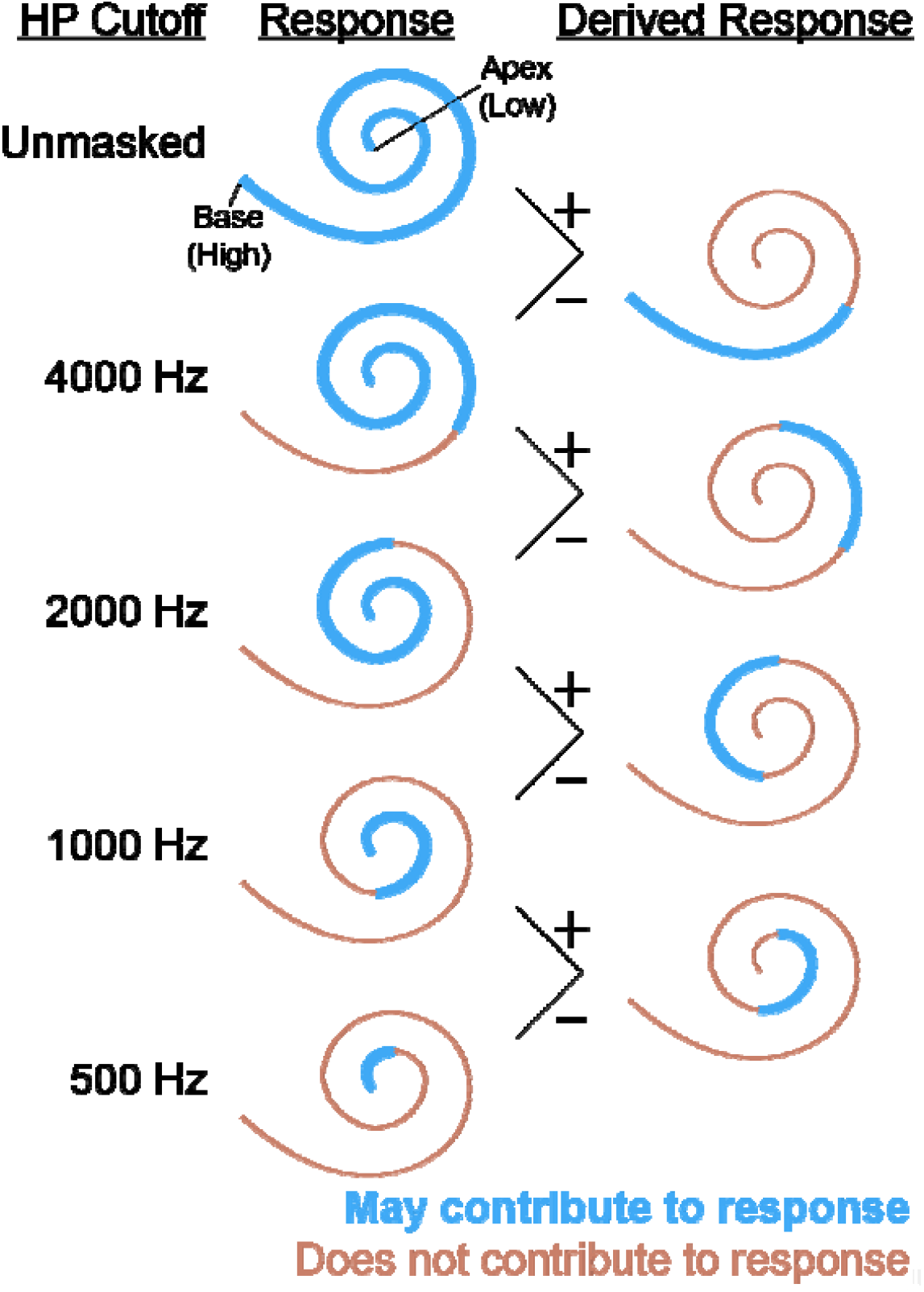
Illustration of the derived-band method for a broadband stimulus. The cutoff frequencies for the high pass noise are shown on the left. Regions of the cochlea which contribute to the response are shown by the thicker blue lines for each HPN masker. The responses can then be subtracted as indicated to produce derived-band responses, with the regions which contribute to these responses shown on the right.

## Methods

### Subjects

The experiment conducted in this study was completed using a protocol approved by the University of Rochester Research Subjects Review Board (No. 3866). We recruited 12 (3 male, 9 female) subjects for this study. All subjects provided informed consent prior to participating in the experiment and were compensated for their time. The mean ± SD age of the subjects was 21 ± 4.1 (range 18–34) years. Subjects reported no neurological disorders or abnormalities. Normal hearing thresholds (≤ 20 dB HL) for all subjects were confirmed with pure tone audiometry from 250 to 8000 Hz at octave frequencies. Due to the large number of conditions (20 HPN cutoffs at 3 stimulus rates) and time needed to test each condition (since the derived response technique halves SNR), recordings were split over 4 sessions. Each session contained 150 minutes of stimuli, providing a total recording time of 600 minutes per subject and 10 minutes per condition. Two subjects completed only two of the four sessions, resulting in only 5 minutes per condition for those subjects.

### pABR generation

The pABR stimuli were constructed as described previously, shown in Fig. 3 [13]. Toneburst trains at five octave-spaced stimulus frequencies (500, 1000, 2000, 4000, and 8000 Hz) were generated with their timing determined by independent random Poisson processes. These toneburst trains were then summed together to produce the stimulus for one ear, and the process was repeated for the other ear. Masking noise (with varying cutoff frequencies) was added to the stimuli (described in more detail below). For both the stimuli and the noise, 60 one-second tokens were generated, and the noise token paired with the stimulus on each trial was randomized.

**Fig. 3.**
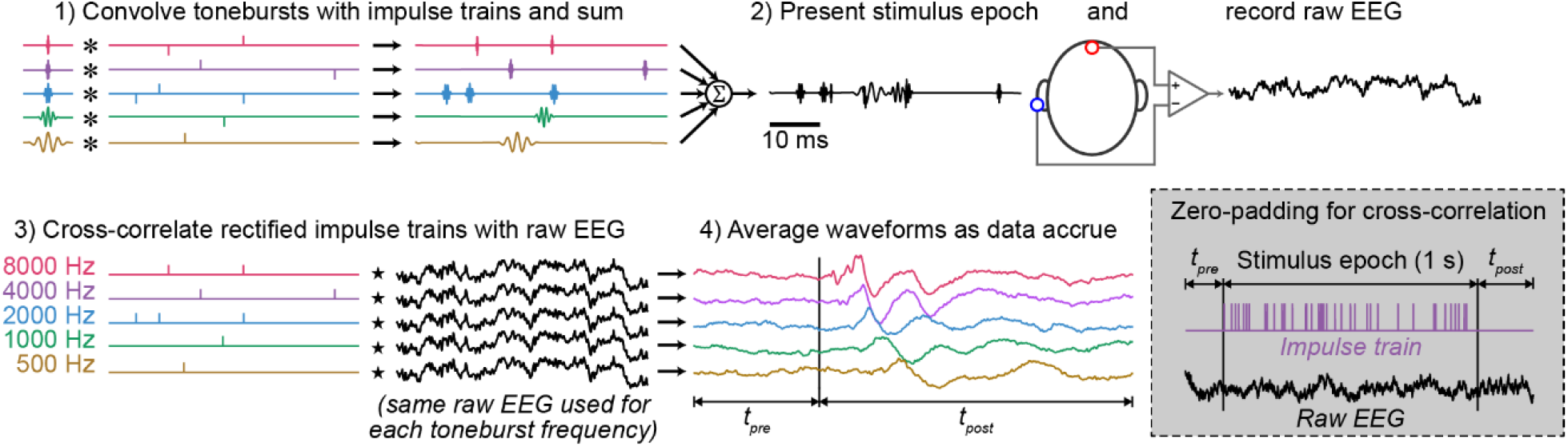
Overview of stimulus generation, recording, and analysis techniques. First, tonebursts of five frequencies are convolved with unique impulse trains and summed together. A random half of the impulses were flipped to produce rarefaction and condensations tonebursts. Next, the stimulus is presented to the subject while recording EEG. Then, the impulse trains used in the stimulus generation step are rectified and cross correlated to determine the response for each stimulus frequency, and responses from different trials are averaged together. Since the cross-correlation is computed in the frequency domain, the impulse train is padded with zeros on either side and the EEG window is expanded to match prior to any calculations to prevent circular artifacts. The process is shown here for one ear, but the other is measured at the same time using differently timed toneburst trains. From Polonenko & Maddox [13]. Licensed CC BY-NC.

Serial stimuli were identical to parallel stimuli, except a single toneburst train was selected to be presented in isolation (i.e., other stimulus frequencies were not included in the stimulus). Only two frequencies (500 Hz and 2000 Hz) were tested for the serial condition due to time constraints during recording. These frequencies were selected because 500 Hz is the lowest commonly used test frequency and expected to have the largest benefit to place specificity from parallel presentation and 2000 Hz is in the middle of the range of commonly tested frequencies, where we expected less difference in place specificity between parallel and serial conditions.

Three stimulus rates (20, 40, and 100 stim/s) were tested to examine the effect of rate on the strength of masking, with different rates presented in a random order. Rates of 20 and 40 stim/s have previously been shown to be optimal to reduce pABR test times [18], while a higher rate of 100 stim/s was expected to provide additional masking and produce more place-specific responses.

### Derived-band responses

Derived-band responses were obtained using methods described in prior studies [15–17]. High pass filtered pink noise was added to both the parallel and serial stimuli, and the cutoff frequency of the high pass filter was varied in half-octave steps. For parallel stimuli, the HPN cutoff frequency ranged from 250 to 16000 Hz, resulting in 13 parallel conditions. For serial stimuli, the cutoff frequency ranged from one octave below to two octaves above the test frequency, resulting in 7 serial conditions. Octave-wide derived responses were obtained by subtracting responses which had the high pass cutoff for the noise filter one octave apart. Throughout this study, we take the convention of denoting the octave-wide response band by the frequency in the center of the two high pass noise cutoffs as determined by the geometric mean (e.g., the derived response band corresponding to the 707 to 1414 Hz region is denoted as 1000 Hz).

### Stimulus presentation and EEG recording

During the experiment, subjects were seated in a comfortable reclining chair in a sound-isolated and electrically shielded room (IAC, North Aurora, IL, USA) and had the option to sleep or watch muted, captioned videos during the experiment. Stimuli were presented using ER-2 insert earphones (ER-2, Etymotic Research, Elk Grove, IL). The toneburst stimuli were presented at 75 dB peSPL. The level from the masking noise was set based on pilot data such that the noise with the lowest cutoff (250 Hz) completely masked the serial responses. Based on pilot data, the masking noise was set to 69 dB SPL, but the level was increased to 72 dB SPL after 6 subjects since responses were not completely masked in all subjects. Masking effects improved, but some subjects still showed responses where they were expected to be masked. In the end, overall results were the same for both masking noise levels, so they are not separated for analysis here.

A python script was used to control the experiment using open-source software [19]. The python script sent both the audio stimuli and triggers through a soundcard (Babyface, RME, Haimhausen, Germany) which sent the triggers to a custom trigger box [20] to be passed to the EEG system for precise timing.

EEG was recorded using BrainVision ActiChamp and EP preamps (Brainvision LLC, Greenboro, SC) at a sampling rate of 10 kHz. Passive Ag/AgCl electrodes were used to record responses from FCz (in the standard 10-20 montage) to right and left earlobe with ground on forehead. The FCz electrode was plugged into a Y-connector to act as the noninverting electrode for both preamps, while the earlobes were used as the inverting electrode.

### Response calculation

The raw EEG recording was bandpass filtered between 30 and 2000 Hz using a causal first order Butterworth filter and notch filtered at odd integer multiples of 60 up to 2500 Hz using causal IIR filters with a bandwidth of 5 Hz. ABRs were calculated as previously described (see Fig. 3). For each subject, the four sessions were concatenated together prior to response calculation. For each stimulus frequency, the impulse train corresponding to the onset of each toneburst of that frequency was rectified and downsampled to match the EEG sampling rate. This was accomplished by taking the index of each impulse, multiplying it by the EEG sampling rate, dividing it by the stimulus sampling rate, and rounding to the nearest integer. The downsampled impulse train was then generated using these indices. All impulses were set to a magnitude of 1 such that rarefaction and condensation responses would average together to cancel stimulus artifact and so the responses had units of volts. These impulse trains (*x*) were then cross correlated with the EEG from that trial (*y*) and divided by the number of stimuli (*n*),

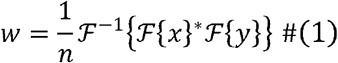

yielding the ABR waveform (*w*), which is mathematically equivalent to averaging but more efficient to compute. For each trial, the impulse trains were zero padded and the EEG was padded with the surrounding waveform prior to moving to response calculation to prevent circular artifacts inherent to working in the frequency domain. For each subject, all trials for a given condition were averaged together using a Bayesian weighting technique, where the weights for a given trial were calculated as the inverse of the variance of the raw EEG for that trial and normalized such that the weights across trials summed to one. This method increases SNR by reducing the contribution of noisy trials to the average response. The responses for serially presented stimuli are calculated in the same way, except there is only one impulse train and one response to be determined.

Both authors picked the wave V peaks together for the grand average un-subtracted responses for later analysis, and TJS picked the peaks of the un-subtracted responses for all subjects. In plots and analyses comparing serial and parallel responses, the parallel response peaks were picked from only the ear from which the serial response was collected. When examining only the parallel condition, responses were averaged across ears prior to peak picking. Individual subjects did not always produce a response for each condition (i.e., at each HPN cutoff for all frequencies and rates). In these cases, the subject was excluded from latency calculations for that condition. In the un-subtracted responses, the latency of the peaks can be used to estimate the place specificity of the response since more basal regions of the cochlea produce responses with shorter latencies. In the derived responses, the magnitude of the response in different bands is used to measure place specificity.

Due to the low SNR of the derived responses, we computed the noise-adjusted standard deviation of the response within the expected response window, *σ*_*R*_, to estimate the magnitude of the response. The latency at which a response was expected was determined by averaging the latencies of the two responses used to generate the derived waveform. We used the latencies from the grand average waveforms to determine the center of the analysis window, because subjects did not always have a response at each condition. The variance of the waveform within an 12 ms time window centered at this latency was calculated which represents the variance of the response plus the noise, 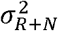. Since the response and noise can be treated as independent random processes, the variance of the response 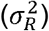 can be estimated by subtracting the variance of the noise 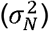 (Equation 2). We estimated the variance of the noise by taking the variance of 40 non-overlapping 12 ms time windows taken from the prestimulus region of the waveform and averaging these variances. This estimate of the noise variance was then subtracted from the 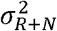,yielding the estimated 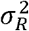. This baseline correction was done separately for each subject and condition. We then took the square root of 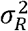 to determine the standard deviation which is more intuitive and has units matching the response waveforms.

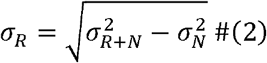

When calculating the response variance to compare the serial and parallel conditions, parallel responses were only taken only from the ear matching that of the serial condition (i.e., responses from both ears were not averaged together), to avoid unfair comparisons due to differences in SNR. The serial and parallel response variances for each stimulus rate and test frequency were then plotted against the derived band center frequency to visualize the excitation pattern.

### Statistical Analysis

All statistical analyses were performed in R v4.3.2 [21]. Linear mixed effects models were fit using the lmer function in the lme4 package [22] from which ANOVAs were performed. All models included the subject identifier as a random effect and stimulus frequency, stimulus rate, HPN cutoff (or derived band), and their interactions as fixed effects. HPN cutoff or derived band was always represented as the difference (in octaves) between the HPN cutoff or derived band center frequency from the test frequency. Separate models were fit for the conditions where data existed to compare serial and parallel responses and to compare the parallel responses to each other. Models comparing serial to parallel responses also included paradigm and its interactions as a fixed effect. All effects were coded as factors in R. The observed variable for the un-subtracted responses was response latency, while the observed variable for the derived responses was response size (*σ*_*R*_ from Equation 2). Serial and parallel response latencies and sizes were compared with post-hoc tests using the emmeans package [23], with adjustments for multiple comparisons using the multivariate t method.

## Results

Our results show parallel presentation improves place specificity for the lower stimulus test frequency (500 Hz) when compared to serial presentation, and there is no difference in place specificity at a higher stimulus frequency (2000 Hz). The improvement seen at the lower test frequency is strongest at higher stimulus rates. Additionally, we show parallel responses at all test frequencies to be place specific.

To demonstrate these results, we first look at grand average responses for the condition where we collected data for both the serial and parallel conditions (a quantitative analysis follows). In Fig. 4a, responses to the 500 Hz toneburst trains presented in serial (darker, dashed lines) or parallel (lighter, solid lines) are shown for the three different stimulus rates (20, 40, and 100 Hz) and all HPN cutoff frequencies where serial responses were collected. At the lowest HPN cutoff frequency, both serial and parallel presentation produce a small and late response. As the HPN cutoff is increased, the responses tend to grow larger and move earlier, as more basal parts of the cochlea are become unmasked. Importantly, the serial responses grow larger and move earlier than the parallel responses – a behavior indicating a comparatively greater basal contribution for serial responses. The difference between the serial and parallel response latencies observed as the HPN cutoff increases is more pronounced at higher stimulus rates. Additionally, responses become smaller with increased stimulus rates, which occurs primarily due to neural adaptation [1, 24]. Fig. 4b shows the responses to the 2000 Hz tonebursts. The serial and parallel responses at that frequency are similar, with no appreciable differences between the amplitudes or latencies at any condition.

**Fig. 4.**
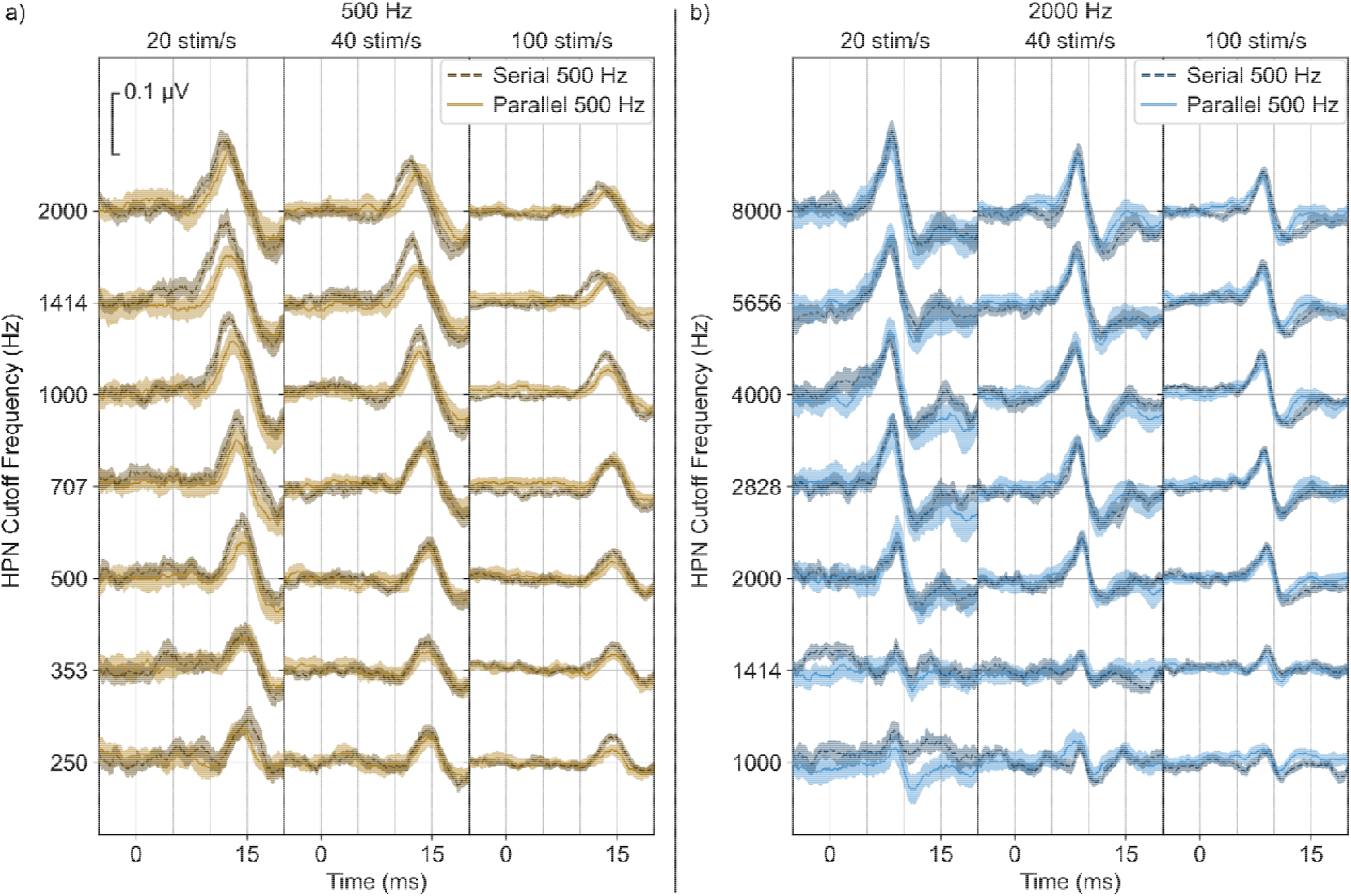
Responses for serial and parallel presentation for the **a)** 500 Hz and **b)** 2000 Hz tonebursts. Within each test frequency, each column represents a different stimulus rate, and each row represents a different HPN cutoff frequency. The HPN cutoff frequency ranged from one octave below to two octaves above the stimulus frequency. Shaded areas indicate ± 1 SEM.

To examine the trends seen in Fig. 4 quantitatively, we plot the wave V latencies for all conditions (Fig. 5), with the serial and parallel comparison in Fig. 5a and all parallel response latencies in Fig. 5b. We additionally used a linear mixed effects model and ANOVA to determine factors which had a significant effect on latency, shown in Table 1. We discuss only the significant variables in the text of the paper, while all effects and interactions are reported in the tables. In the 500 Hz condition, parallel responses latencies at the two highest HPN cutoff frequencies are later than the serial response latencies, with greater separation at higher stimulus rates. The main effect of stimulus rate and the interaction of rate with HPN cutoff frequency indicates the change in latency with stimulus rate is significant and only present when less masking is present, while the effect of paradigm indicates the differences between serial and parallel latencies are significant. The difference between serial and parallel latencies is only present at higher HPN cutoff frequencies, as indicated by the significant effect of HPN cutoff frequency and the interaction between paradigm and HPN cutoff frequency in the LME model and confirmed through post-hoc testing. For the 2000 Hz condition, we see no difference between the serial and parallel latencies at any HPN cutoff for all stimulus rates. The main effect of stimulus frequency reflects the lower latency of the 2000 Hz responses, while the interactions between stimulus frequency and the other variables indicate that latency differences were only present for the 500 Hz responses.

**Table 1.** Type-III ANOVA on the LME model to compare serial and parallel response latencies.

| Variable | DF<sub>Num</sub> | DF<sub>Den</sub> | F value | p |  |
| --- | --- | --- | --- | --- | --- |
| Paradigm | 1 | 861 | 3.86 | 0.0497 | * |
| Rate | 2 | 858 | 21.2 | $1.05 \times 10^{-9}$ | *** |
| Stimulus Frequency | 1 | 858 | 8860 | $<2.2 \times 10^{-16}$ | *** |
| HPN Cutoff Frequency | 6 | 858 | 41.6 | $<2.2 \times 10^{-16}$ | *** |
| Paradigm: Rate | 2 | 858 | 0.321 | 0.72 |  |
| Paradigm: Stimulus Frequency | 1 | 858 | 36.5 | $2.27 \times 10^{-9}$ | *** |
| Rate: Stimulus Frequency | 2 | 858 | 1.90 | 0.150 |  |
| Paradigm: HPN Cutoff Frequency | 6 | 858 | 5.95 | $4.24 \times 10^{-6}$ | *** |
| Rate: HPN Cutoff Frequency | 12 | 858 | 2.24 | $8.81 \times 10^{-3}$ | ** |
| Stimulus Frequency: HPN Cutoff Frequency | 6 | 858 | 10.0 | $1.05 \times 10^{-10}$ | *** |
| Paradigm: Rate: Stimulus Frequency | 2 | 858 | 1.76 | 0.173 |  |
| Paradigm: Rate: HPN Cutoff Frequency | 12 | 858 | 0.237 | 1.00 |  |
| Paradigm: Stimulus Frequency: HPN Cutoff Frequency | 6 | 858 | 6.52 | $9.59 \times 10^{-7}$ | *** |
| Rate: Stimulus Frequency: HPN Cutoff Frequency | 12 | 858 | 0.239 | 1.00 |  |
| Paradigm: Rate: Stimulus Frequency: HPN Cutoff Frequency | 12 | 858 | 0.520 | 1.00 |  |

**Fig. 5.**
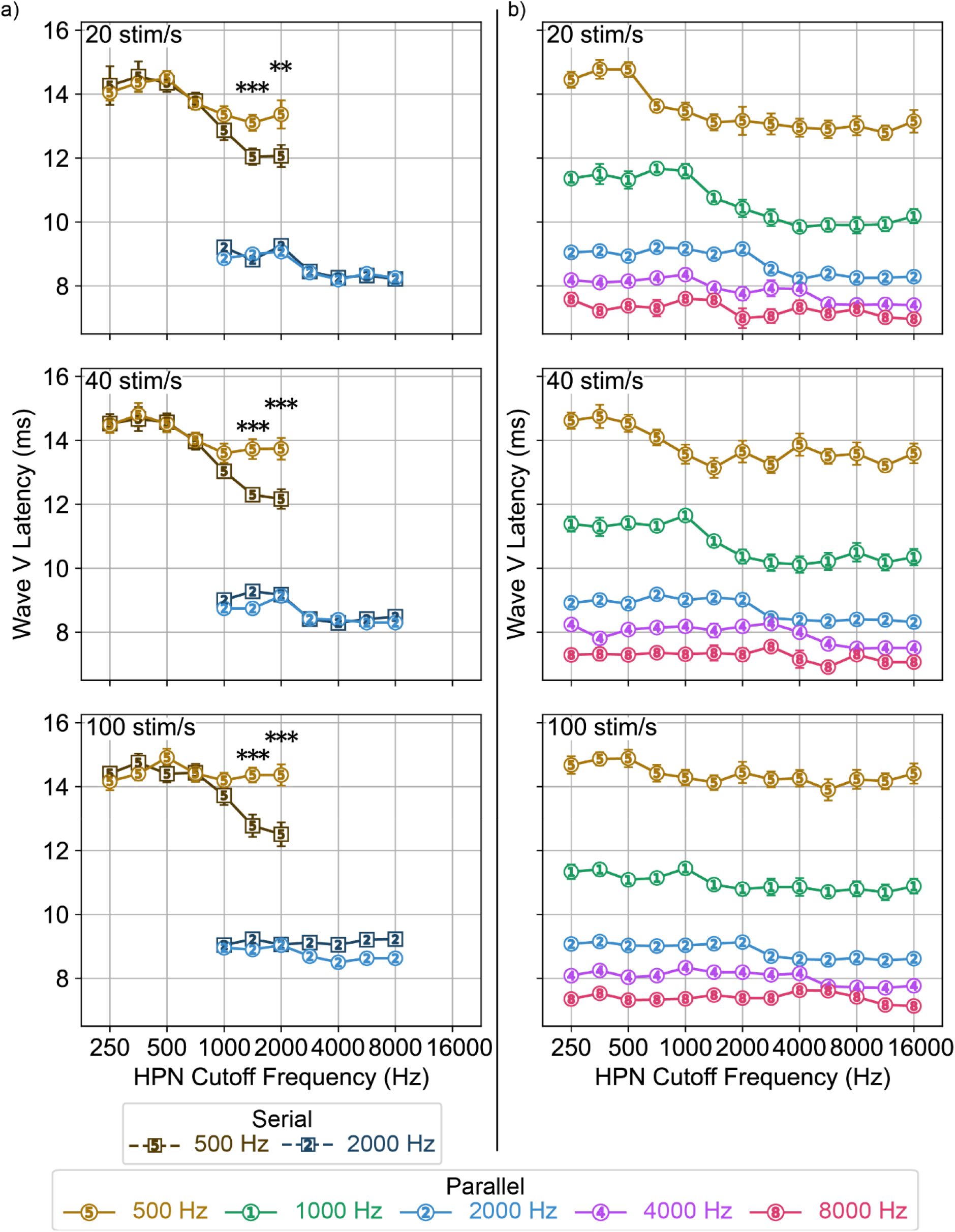
Wave V response latencies across HPN cutoff frequency. **a)** Latencies for the serial and parallel responses to the 500 and 2000 Hz stimuli. Responses to the 500 Hz stimuli had later latencies with parallel presentation than serial at the two highest HPN cutoff frequencies for all rates (** p<0.01, *** p<0.001) **b)** Latencies for responses to all parallel stimulus frequencies. Error bars indicate ± 1 SEM.

There is also clear variation in the parallel response latencies shown in Fig. 5b. Table 2 shows the results of the LME model. The main effect of stimulus frequency represents the latency difference for responses to different frequencies. Responses to higher stimulus frequencies had more consistent latencies across both HPN cutoff and stimulus rate, supported by the effects of rate and HPN cutoff and their interactions with stimulus frequency. The interaction between rate and HPN cutoff frequency indicates a that stimulus rate had an effect on response latency, likely reflecting the later latencies observed at higher HPN cutoff frequencies at the higher stimulus rates in Fig. 5b.

**Table 2.** Type-III ANOVA on the LME model comparing parallel response latencies.

| Variable | DF<sub>Num</sub> | DF<sub>Den</sub> | F value | p |  |
| --- | --- | --- | --- | --- | --- |
| Rate | 2 | 1209 | 30.5 | $1.20 \times 10^{-13}$ | *** |
| Stimulus Frequency | 4 | 1209 | 6320 | $<2.2 \times 10^{-16}$ | *** |
| HPN Cutoff Frequency | 3 | 1209 | 181 | $<2.2 \times 10^{-16}$ | *** |
| Rate: Stimulus Frequency | 8 | 1209 | 3.45 | $6.61 \times 10^{-4}$ | *** |
| Rate: HPN Cutoff Frequency | 6 | 1209 | 8.39 | $6.20 \times 10^{-9}$ | *** |
| Stimulus Frequency: HPN Cutoff Frequency | 11 | 1209 | 7.18 | $6.06 \times 10^{-12}$ | *** |
| Rate: Stimulus Frequency: HPN Cutoff Frequency | 22 | 1209 | 1.81 | 0.0121 | * |

To more closely examine which frequency bands of the auditory system contributed to the responses, we calculated octave-wide derived responses for all conditions. Here, we examine the presence or absence of a response in each derived band to evaluate response spread. Fig. 6 shows the serial and parallel derived responses for 500 (Fig. 6a) and 2000 Hz (Fig. 6b). Qualitatively, the 500 Hz serial responses are larger and present in more response bands than the 500 Hz parallel responses, with greater differences at the higher rates. There is again little difference between the serial and parallel 2000 Hz responses.

**Fig. 6.**
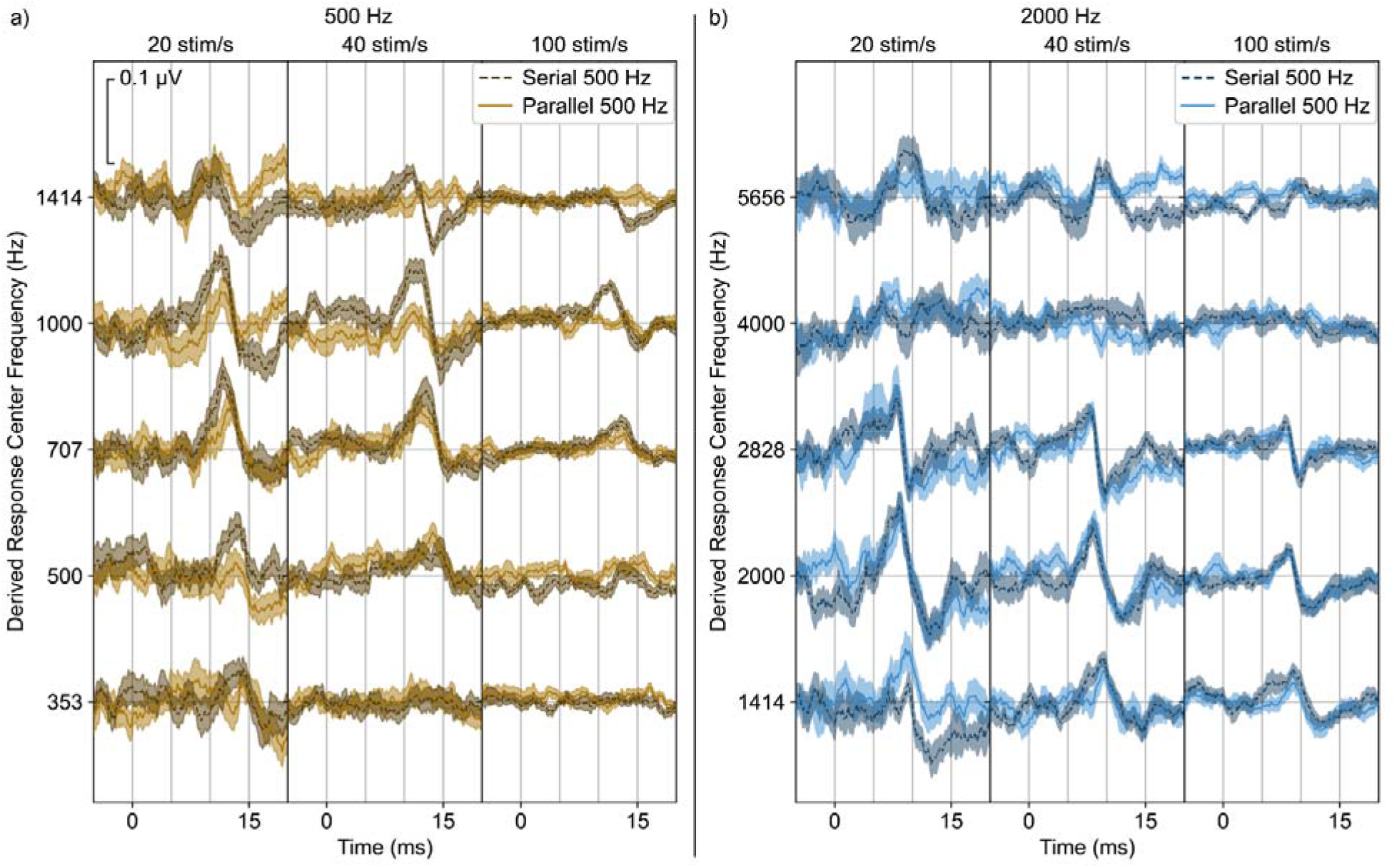
Derived responses for serial (darker, dashed line) and parallel (lighter, solid line) presentation for **a)** 500 Hz tonebursts and **b)** 2000 Hz tonebursts. Shaded areas indicate ± 1 SEM.

To better characterize the responses in Fig. 6, we estimated the response size (as described in Methods – response size could be negative due to bias correction), shown in Fig. 7. With an estimate of the response size, we again fit an LME model to the data to compare the serial and parallel responses shown in Fig. 7a. The results of this model are shown in Table 3. The effect of derived band indicates the amplitude of the response changed across derived response band. Stimulus rate also influenced the amplitude of the responses, likely reflecting the reduction in response size with increasing stimulus rate. The effect of paradigm and its interaction with stimulus frequency and derived band indicates parallel presentation affected response size differently depending on the derived band and stimulus frequency. This reflects the 500 Hz response becoming narrower and shifting toward lower derived bands for parallel responses while the 2000 Hz responses remained similar in shape and location. The interaction of stimulus frequency and derived band reflects the different shapes of the response patterns for the two stimulus frequencies, while the interaction of rate, stimulus frequency, and derived band shows that rate affects the shape of the responses more for one frequency than the other. Specifically, parallel presentation reduced the size of the 500 Hz responses one octave above the test frequency. This effect was present at all rates, with the greatest difference at 40 stim/s. There was also a significant difference between the size of the serial and parallel 2000 Hz response one and a half octaves above the test frequency at a rate of 40 stim/s – we suspect this is a type I error because it is not predicted by our understanding of the system or consistent with the other trends in the data, but report it here for completeness.

**Table 3.**
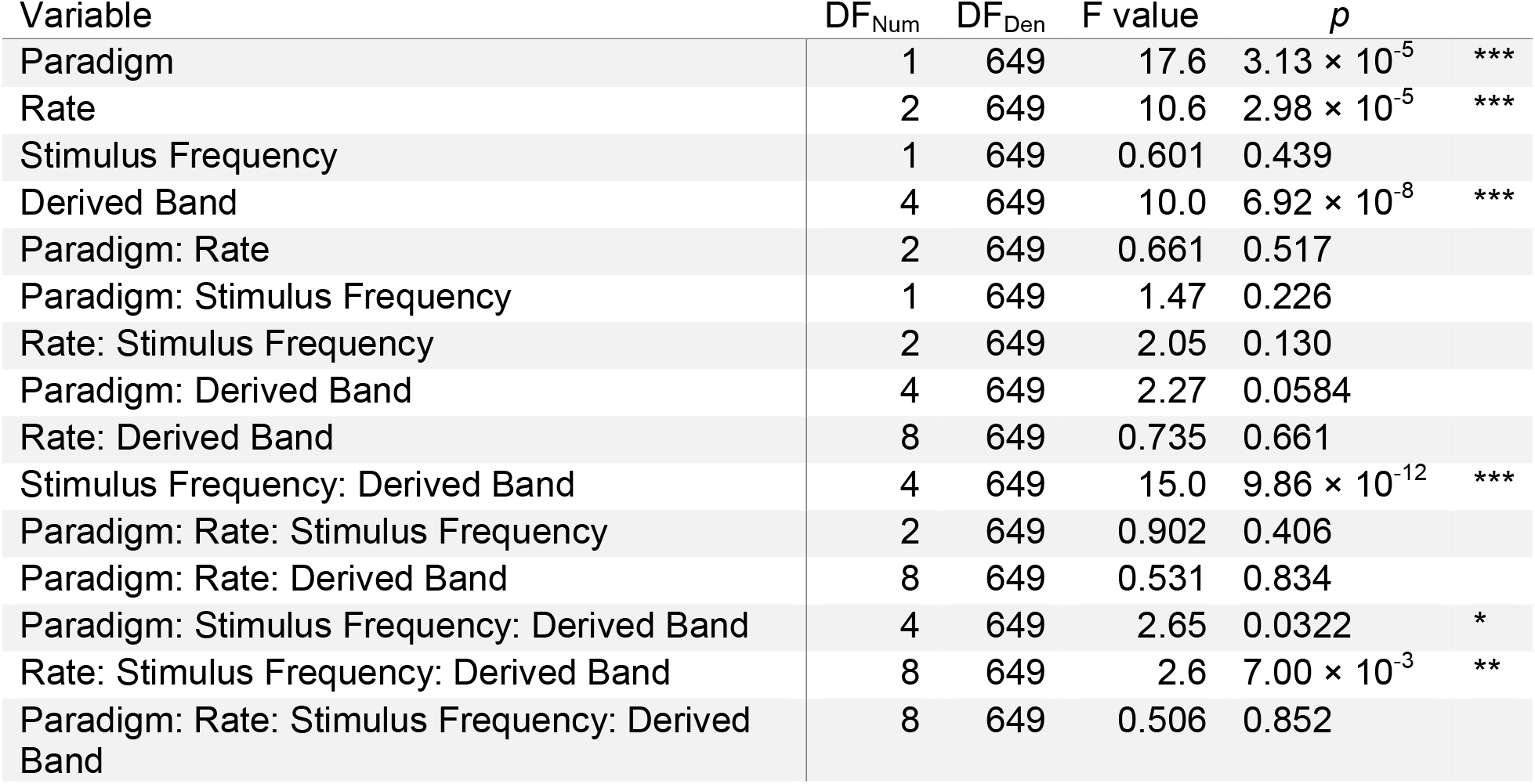
Type-III ANOVA on the LME model comparing serial and parallel response size.

**Fig. 7.**
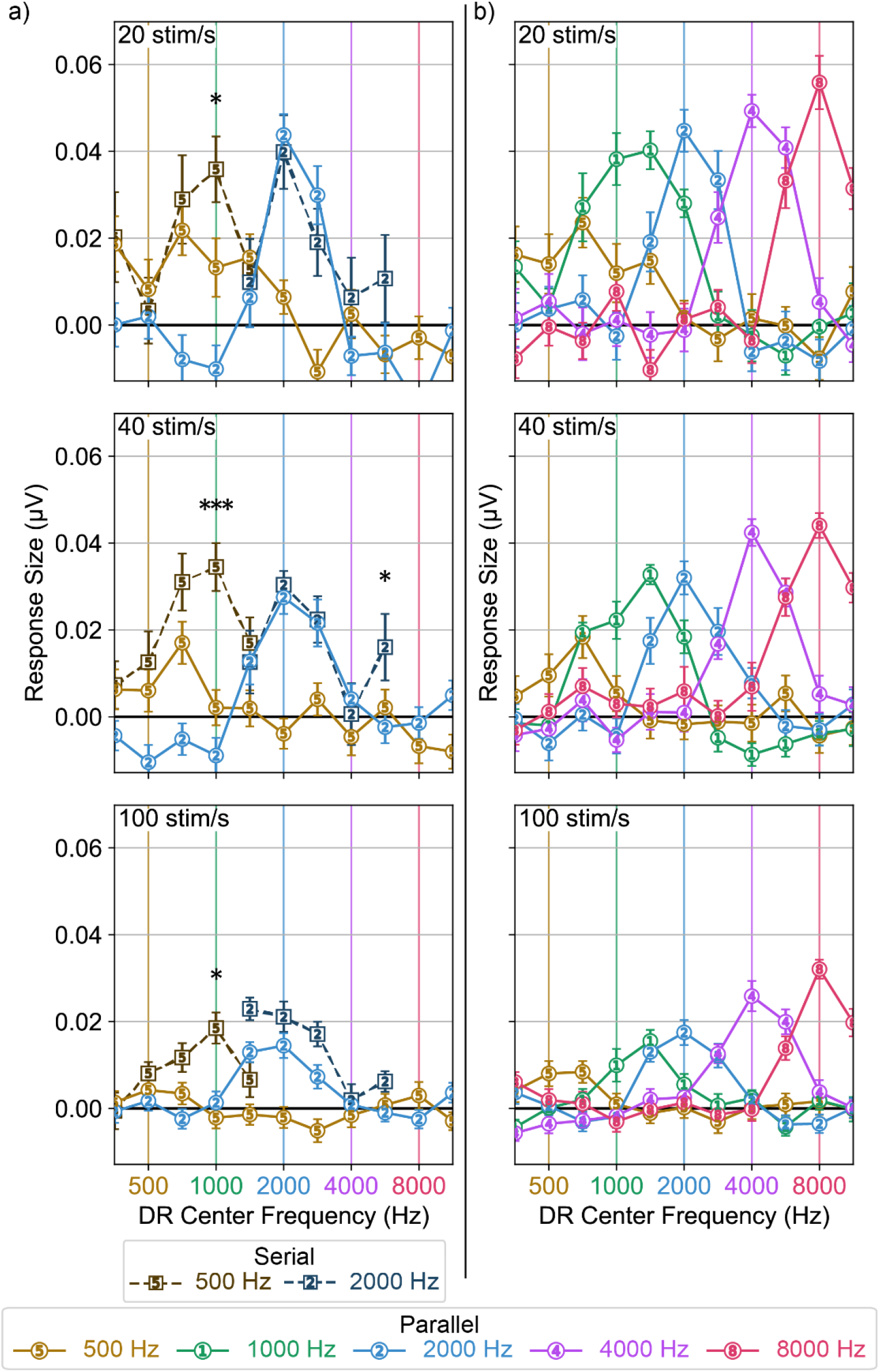
Response size within each derived response band. **a)** Serial and parallel response sizes for the 500 and 2000 Hz stimuli. Parallel presentation reduced response size in the derived band one octave above the stimulus frequency for the 500 Hz responses at all rates (* p<0.05, ** p<0.01, *** p<0.001), **b)** Response sizes for all parallel stimulus frequencies. Response size was estimated as the standard deviation of the response in the expected response window, baselined by that of the prestimulus response. Error bars indicate ± 1 SEM.

We again fit a separate model to compare the parallel responses (Fig. 7b) with one another, shown in Table 4. The main effects stimulus rate and stimulus frequency reflect the differences in response size as these variables change, while the effect of derived band again reflects the varying response size across derived response bands. There were interactions between stimulus rate and derived band as well as between stimulus frequency and derived band, indicating that stimulus rate and frequency both affected the shape of the response across frequency.

**Table 4.** Type-III ANOVA on the LME model comparing parallel response size.

| Variable | DF<sub>Num</sub> | DF<sub>Den</sub> | F value | p |  |
| --- | --- | --- | --- | --- | --- |
| Rate | 2 | 979 | 31.5 | $4.02 \times 10^{-14}$ | *** |
| Stimulus Frequency | 4 | 979 | 17.6 | $5.37 \times 10^{-14}$ | *** |
| Derived Band | 6 | 979 | 109 | $<2.2 \times 10^{-16}$ | *** |
| Rate: Stimulus Frequency | 8 | 979 | 1.74 | 0.0857 |  |
| Rate: Derived Band | 12 | 979 | 5.66 | $1.78 \times 10^{-9}$ | *** |
| Stimulus Frequency: Derived Band | 19 | 979 | 6.34 | $1.16 \times 10^{-15}$ | *** |
| Rate: Stimulus Frequency: Derived Band | 38 | 979 | 1.17 | 0.229 |  |

## Discussion

In this study, we investigated how parallel presentation affects place specificity of ABRs at different stimulus rates and frequencies. We found that parallel presentation improves place specificity for the 500 Hz stimulus frequency with more improvement at higher rates, as evidenced by the increasing latency difference between serial and parallel presentation in the un-subtracted responses (Fig. 5a) and greater difference in response size in derived response bands above the stimulus frequency (Fig. 7a). Parallel presentation produced responses which were at least as place specific as responses from serial presentation for the 2000 Hz stimulus frequency at all rates, with a possible improvement at 40 stim/s (Fig. 7a), although this improvement was only marginally significant and not seen in the response latencies (Fig. 5a). All parallel stimuli produced place-specific responses, with response patterns typically centered on their test frequencies and response strength diminishing quickly when moving away from test frequency.

Here, we were able to examine place specificity of the pABR in human subjects, supporting our findings from a previous modeling study examining the same question [14]. While we were only able to compare serial and parallel presentation for two stimulus frequencies, all parallel stimulus frequencies were able to be compared. Comparisons were complicated by the noise in the responses, particularly in the derived responses which had substantially worse SNR than the un-subtracted responses due to smaller responses and double the noise (both effects resulting from two similar measurements with independent noise being subtracted). This prevented us from using a direct measurement of wave V amplitude, so we instead relied on the noise-corrected standard deviation in the response window.

Even at the lowest HPN cutoff frequency, responses were not completely masked, so there were likely contributions to the response from higher frequency regions than expected in each derived band due to incomplete masking. Masking was improved for the latter half of the subjects, but the masking of responses remained incomplete. During analysis, we qualitatively compared the grand averages using only the first half and latter half of subjects and saw no difference between the two groups.

This study closely followed the experimental design of previous studies which that investigated place specificity of EEG responses [16, 17]. As in these prior studies, we find the 2000 Hz serial response to be place specific. Unlike the prior studies, we see somewhat poor place specificity for the 500 Hz serial responses, with the greatest proportion of the response generated by the frequency band centered an octave above the stimulus frequency. This shift could partially be explained by the incomplete masking, which allows for some contributions from higher frequency bands.

Our finding of improved place specificity for the 500 Hz responses supports the findings of our previous study which used computational models to investigate place specificity of the pABR [14]. However, the modeling study suggested improvements would be present for almost all stimulus frequencies (depending on the stimulus rate and level), which is not supported by the present study. Several factors may contribute to this difference. First, the modeled responses had no noise, allowing for observations of differences that may be hidden in recorded data.

Secondly, models are a simplification of the true system, requiring compromises to make building and using the model feasible and representing only our best understanding of the system being modeled. There may be key factors which are missing from that model that would reduce the differences in the model findings and EEG findings. Lastly, we based our analysis here on the latency and derived amplitude of ABR wave V. In recorded data, this is sensible since wave V is the most robust component of the ABR and has the greatest clinical relevance. However, in the modeling study, we focused on auditory nerve responses, which correspond to wave I, since models of the auditory periphery are more advanced than models of the neural generators for wave V. Responses may be altered as they move from the auditory nerve to the brainstem in a way that would affect our estimates of place specificity. For example, our analyses relied, in part, on the latency difference expected from different response bands. However, some studies have reported that octopus cells in the cochlear nucleus can reduce this frequency-dependent latency difference [25–27], which would make responses from different frequency bands more similar.

Based on our past work which found rates of 20 or 40 stim/s to provide the greatest improvement in test times for parallel versus serial stimulation, and this study which finds the greatest improvements in place specificity for the 500 Hz stimulus at higher rates, a stimulus rate of 40 stim/s seems to be optimal for the pABR, at least in adult subjects. When using the stimulus rate of 40 stim/s, the pABR is as place specific or better than serial methods while also producing responses in less time. In addition to the faster data collection times, threshold estimates may also improve since notched-noise masking has been shown to reduce differences between thresholds estimated with ABRs and those measured behaviorally [8].

One downside of the pABR is that its speed is limited by the slowest of the ten responses being simultaneously recorded. The slowest response to measure is 500 Hz in the vast majority of cases [13]. We have previously reasoned that slower parallel than serial 500 Hz response acquisition is because of the spurious basal contributions of the serial response being masked out. A closer inspection of Fig. 7b, however, suggests that at 20 and 40 stim/s, the 1000 Hz stimulus doesn’t just mask unwanted 500 Hz responses, but may be spreading apically in a way that impedes the 500 Hz response from the true 500 Hz portion of the tonotopic axis. This finding would suggest that increasing the amplitude of the 500 Hz stimuli relative to the other frequencies being presented in parallel may speed collection of the rate-limiting 500 Hz responses (while maintaining place specificity), thus speeding pABR collection overall.

## Conclusions

The parallel presentation used in the pABR paradigm provides place specific responses, and can improve place specificity relative to serial presentation for low stimulus frequencies, especially at higher stimulus rates. This benefit is due to masking effects inherent to the stimulus construction, a design which speeds data collection, in contrast to other masking techniques which prolong it. Based on our previous work examining stimulus rates for faster test times and this study which examined the effect of different stimulus rates on place specificity, we find that a stimulus rate of 40 stim/s is optimal for the pABR paradigm.

## Acknowledgements

The authors thank Yathida Melody Anankul for assistance with data collection. This work was supported by NIH grant R01DC017962 awarded to RKM.

## References

1. Burkard RF, Eggermont JJ, Don M (2007) Auditory evoked potentials: basic principles and clinical application. Lippincott Williams & Wilkins

2. Robles L, Ruggero MA (2001) Mechanics of the Mammalian Cochlea. Physiological Reviews 81:1305–1352. 10.1152/physrev.2001.81.3.1305

3. Russell IJ, Nilsen KE (1997) The location of the cochlear amplifier: Spatial representation of a single tone on the guinea pig basilar membrane. Proceedings of the National Academy of Sciences 94:2660–2664. 10.1073/pnas.94.6.2660

4. Pickles JO (2012) Introduction to the Physiology of Hearing. BRILL, Bradford, UNITED KINGDOM

5. Purdy SC, Abbas PJ (2002) ABR Thresholds to Tonebursts Gated with Blackman and Linear Windows in Adults with High-Frequency Sensorineural Hearing Loss. Ear and Hearing 23:358–368. 10.1097/00003446-200208000-00011

6. Picton TW (1978) The strategy of evoked potential audiometry. Early diagnosis of hearing loss 297–307

7. Stapells D, Gravel J, Martin B (1995) Thresholds for Auditory Brain Stem Responses to Tones in Notched Noise from Infants and Young Children with Normal Hearing or Sensorineural Hearing Loss. Ear and hearing 16:361–71. 10.1097/00003446-199508000-00003

8. Hood LJ (1998) Clinical applications of the auditory brainstem response. Singular

9. Picton T, Ouellette J, Hamel G, Durieux-Smith A (1979) Brain Stem Evoked Potentials to Tone Pips in Notched Noise. The Journal of otolaryngology 8:289–314

10. Purdy SC, Houghton JM, Keith WJ, Greville KA (1989) Frequency-Specific Auditory Brainstem Responses: Effective Masking Levels and Relationship to Behavioural Thresholds in Normal Hearing Adults. Audiology 28:82–91. 10.3109/00206098909081613

11. Hall JW (2007) New handbook of auditory evoked responses. Pearson

12. BC Early Hearing Program (2012) BC Early Hearing Program. Audiology Assessment Protocol Verson 4:18

13. Polonenko MJ, Maddox RK (2019) The Parallel Auditory Brainstem Response. Trends in Hearing 23:2331216519871395. 10.1177/2331216519871395

14. Stoll TJ, Maddox RK (2023) Enhanced Place Specificity of the Parallel Auditory Brainstem Response: A Modeling Study. Trends in Hearing 27:23312165231205719. 10.1177/23312165231205719

15. Oates P, Stapells D (1997) Frequency specificity of the human auditory brainstem and middle latency responses to brief tones. I. High-pass noise masking. The Journal of the Acoustical Society of America 102:3597–3608. 10.1121/1.420148

16. Oates P, Stapells D (1997) Frequency specificity of the human auditory brainstem and middle latency responses to brief tones. II. Derived response analyses. The Journal of the Acoustical Society of America 102:3609–3619. 10.1121/1.420400

17. Herdman AT, Picton TW, Stapells DR (2002) Place specificity of multiple auditory steady-state responses. The Journal of the Acoustical Society of America 112:1569–1582. 10.1121/1.1506367

18. Polonenko MJ, Maddox RK (2022) Optimizing Parameters for Using the Parallel Auditory Brainstem Response to Quickly Estimate Hearing Thresholds. Ear and hearing 43:646–658. 10.1097/aud.0000000000001128

19. Larson E, McCloy D, Maddox R, Pospisil D (2015) expyfun: Python experimental paradigm functions, version 2.0.0. 10.5281/zenodo.11640

20. Maddox RK (2020) S/Plitter: hardware and firmware for converting digital audio to TTL triggers. 10.5281/zenodo.10802517

21. R Core Team (2023) R: A Language and Environment for Statistical Computing. R Foundation for Statistical Computing, Vienna, Austria

22. Bates D, Mächler M, Bolker B, Walker S (2015) Fitting Linear Mixed-Effects Models Using lme4. Journal of Statistical Software 67:1–48. 10.18637/jss.v067.i01

23. Lenth RV (2023) emmeans: Estimated Marginal Means, aka Least-Squares Means

24. Picton TW (2010) Human Auditory Evoked Potentials. Plural Publishing, Inc., San Diego, UNITED STATES

25. Spencer MJ, Meffin H, Burkitt AN, Grayden DB (2018) Compensation for Traveling Wave Delay Through Selection of Dendritic Delays Using Spike-Timing-Dependent Plasticity in a Model of the Auditory Brainstem. Frontiers in Computational Neuroscience 12:. 10.3389/fncom.2018.00036

26. Spencer M, Grayden D, Bruce I, Meffin H, Burkitt A (2012) An investigation of dendritic delay in octopus cells of the mammalian cochlear nucleus. Frontiers in Computational Neuroscience 6:. 10.3389/fncom.2012.00083

27. Golding NL, Ferragamo MJ, Oertel D (1999) Role of Intrinsic Conductances Underlying Responses to Transients in Octopus Cells of the Cochlear Nucleus. J Neurosci 19:2897–2905. 10.1523/JNEUROSCI.19-08-02897.1999

